# Investigating peripubertal stress and exogenous corticosterone as endogenous stimulators of brain metabolism

**DOI:** 10.1101/2020.07.14.203067

**Authors:** Nathalie Just

**Affiliations:** INRAe Centre Val de Loire. France; Animal Imaging and Technology Core. Ecole Polytechnique Fédérale de Lausanne. Lausanne. Switzerland

**Keywords:** peripubertal stress, corticosterone, individual stress, paternal stress, fMRS, septal

## Abstract

Substantial research on the association between early-life stress and its long-lasting impact on lifetime mental health has been performed revealing that early-life environmental adversity strongly regulates brain function. Alterations of gene expression and behavior in the off-springs of paternally stressed rats were also revealed. However, the precise mechanisms underlying these changes remain poorly understood. Here, an improved characterization of these processes from investigations of the functional metabolism of animal models exposed to peripubertal stress (PS) is proposed. The ultimate goal of this study was to bring forward functional Magnetic Resonance Spectroscopy (fMRS) as a technique of interest for a better understanding of brain areas by endogenous stimulators such as stress. The present study evaluated, compared and classified effects of individual PS (iPS) and paternal PS (pPS) under corticosterone (CORT) challenge in the septal areas of adult rats. Acute stress was simulated by injection of CORT and metabolic concentration changes were analyzed as a function of time. Evaluation of Glucose and Lactate concentration changes allowed the classification of groups of rats using a Glc to Lac index. Moreover, metabolic responses of control rats (CC) and of pPS x iPS rats (SS) were similar while responses in pPS (SC) and iPS (CS) differed, revealing differential adaption of energetic metabolism and of glutamatergic neurotransmission. Findings have crucial interest for understanding the metabolic mechanisms underlying altered functional connectivity and neuronal plasticity in septal areas inducing increased aggressivity in early-life stressed rats.

## INTRODUCTION

The past decade has seen spectacular advances in understanding the brain mechanisms underlying the psychopathology of various disorders such as stress. anxiety. depression and fear (1, 2, 3). These findings are fundamental to disentangle the links between stress disorders and psychiatric diseases. In pathologies such as schizophrenia (SZ). the elevated risk of aggressive behavior has been consistently reported. However, mechanisms underlying it e.g. genetic factors, development of neurocognitive impairments, substance abuse, childhood maltreatment or stress, are difficult to identify (4).

To date, the *in-vivo* characterization of the physiological and neural brain mechanisms induced by social stress has been mainly performed with behavioral, electroencephalographic (EEG) measurements, functional magnetic resonance imaging (fMRI) and/or positron emission tomography (PET) both in clinical and experimental research (5, 6). Meanwhile. alterations of brain energetics due to prolonged situations of stress have also been measured thereby demonstrating a wealth of results allowing increased understanding of the underpinnings of stress processes and consequences (7, 8).

The brain responds to stressful conditions by the release of corticosteroids mediating neuronal action in a complex time-dependent and region-dependent fashion. Activity is enhanced in areas controlling emotions and behavioral strategies while cognitive functions also become activated allowing dedicated brain areas to store information for future use. Upon repeated stress exposure, these normal responses may become compromised. Several investigations already associated early life stress (ELS) and the development of adult psychiatric disorders (9). Notably, hormonal changes due to exposure of the developing brain to severe or prolonged stress may affect physiological, metabolic and endocrine functions. For example, rats exposed to repeated restraint at adolescence have higher corticosterone (CORT) plasma levels. In these rats, it takes longer for CORT levels to come back to normal levels at adulthood after stressful episodes. One hypothesis to explain these phenomena may be that CORT is necessary to mobilize more energy in response to stress at adolescence, thus compensating for higher energetic demands at this period of age. However, other studies argue that the energetic reserves are not sufficient for compensatory mechanisms. Unfortunately, prolonged exposure to CORT also induces prolonged metabolic and adaptive effects as well as cognitive alterations. It is therefore fundamental to understand the physiological and behavioral mechanisms underlying the alteration of hormonal response by stress during prepubertal stages.

In addition, the behavior of individuals may be changed for life. Excessive fear, depression, addiction, eating disorders and violence can develop as a consequence of ELS. In a previous study (7), adult rats subjected to short unpredictable episodes of peripubertal stress (PS) during their childhood (iPS) or during the childhood of their fathers (pPS) demonstrated increased aggressive and anxiety-like behaviors linked to specific changes in septal metabolism assessed with proton MR spectroscopy (^1^H-MRS). Under basal conditions, increased levels of glucose (Glc) and lactate (Lac) were found in all groups of rats relative to controls (CC) (7). The rising endogenous concentrations of glucocorticoids due to increased stress levels increased circulating levels of Glc (10). Moreover, Lac is known as an additional source of energy under stressful conditions (10), which production and release result from brain Glc utilization. Interestingly, basal glucose levels were substantially higher in pPS rats without substantial changes in glutamate (Glu) and glutamine (Gln) mean concentrations relative to controls while the Glu to-Gln-to GABA ratios remained rather constant across groups of rats suggesting adaptive mechanisms for the maintenance of neurotransmission in pPS and iPS animals. Altogether, these results supported increased oxidative metabolism as a result of PS (pPS and iPS without distinction). In addition. these findings supported the view of stress as a modulator of brain neuro-energetics (10).

Proton functional MR Spectroscopy (^1^H-fMRS) allows for the quantification of metabolite concentration changes as a function of time during focal brain activity. This modality has been successfully conducted during visual or cognitive tasks in the healthy brain in clinical and experimental research (11, 12, 13, 14) and is rapidly becoming a complementary technique to fMRI in studies related to pain (15), SZ (16) or stress (8). Both clinical and experimental ^1^H-fMRS studies supported increased oxidative metabolism in response to an external stimulus (13, 14). During stress, neurons are exposed to waves of hormones, which can alter their activity over the course of several hours. Glucocorticoids and their receptors play an important role in the control of neuronal activity (17). Moreover, they participate in the control of whole-body homeostasis (10) as well as in adaption and response of the organism to stress. Exogenous CORT injection is a widely used preclinical model serving to mimic mental disorders and depression at adulthood after repeated episodes of stress during adolescence. Nevertheless, neurobiological effects of CORT remain poorly understood. In this context, using ^1^H-fMRS in conjunction with a pharmacological paradigm such as CORT challenge appears relevant.

Here, ^1^H-fMRS was conducted assuming that pPS, iPS and CORT act as endogenous stimulators of brain activity. The neurochemical consequences of exogenous CORT on septal metabolism in paternal PS (pPS) and/or in individual PS (iPS) rats was also examined.

## Materials and Methods

### Animals

All studies were performed in accordance with the local ethics committee and met the guidelines of the Swiss Cantonal Veterinary office Committee for Animal Experimentation.

Male and Female Wistar Han rats were purchased from Charles River laboratories (Charles River, l’Arbresle, France). Animals were maintained under controlled conditions (12h light/dark. 22±2 °C) and were given food and water *ad libitum*. Male Wistar rats were peripubertally stressed following previously described protocols (18, 19). Briefly, male Wistar Han offsprings were born from 24 male Wistar Han either sub-chronically exposed to stress (predator odor and open elevated spaces) during peripuberty (7 days of stress across the P28-P42 period) or from 24 other male Wistar Han left undisturbed. After randomization processes, 24 male offsprings were further subdivided and exposed to peripubertal stress under the same conditions exposed earlier or left undisturbed (**Fig 1**).

**Figure 1:**
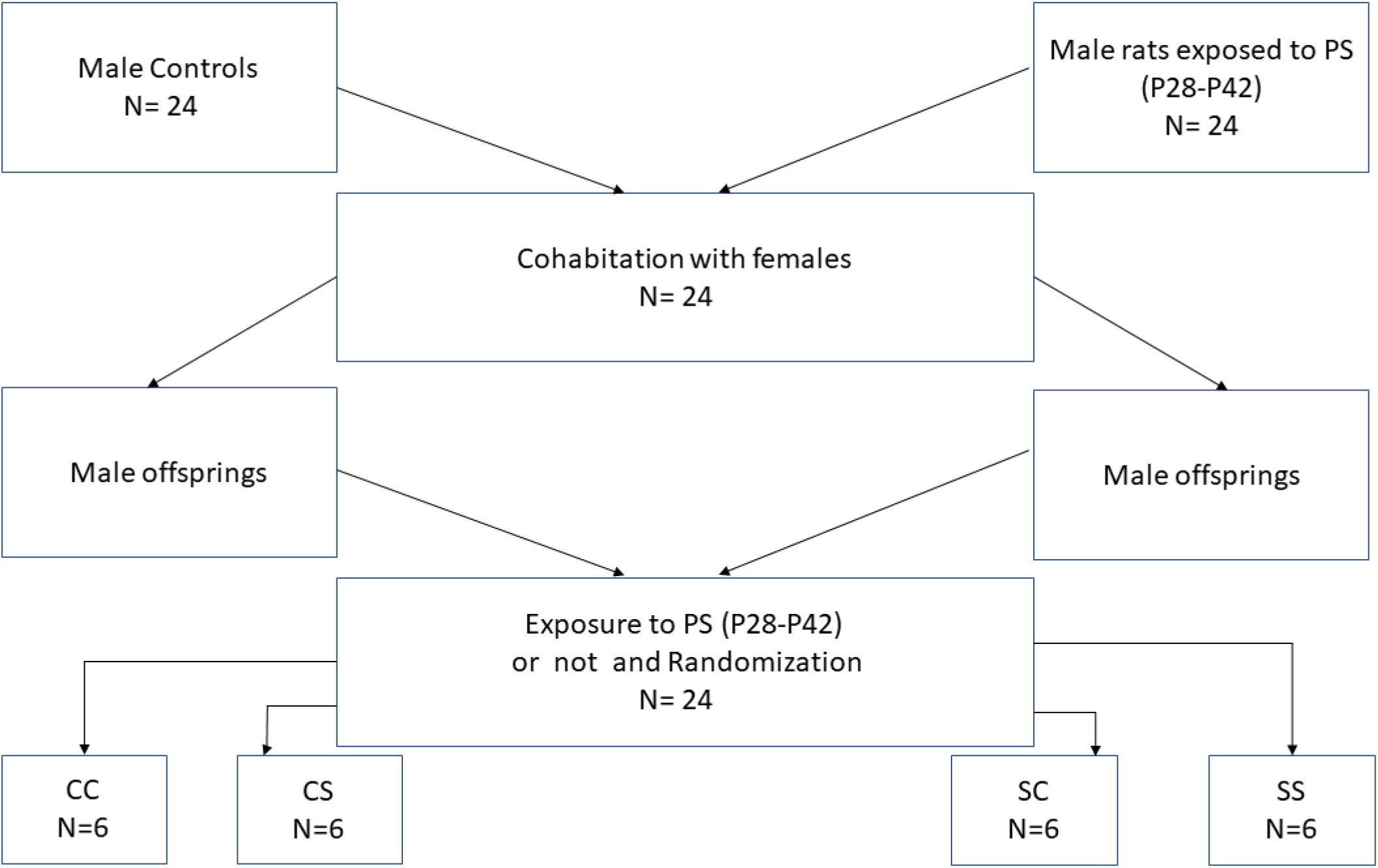
Schematic representation of the procedures leading to 4 groups of n=6 rats: CC, CS, SC and SS. C= Control, S= Stress. The first letter represents the condition of fathers (affected by PS or not), the second letter represents the condition of male offsprings (affected by PS or not).

For further MRI and MRS acquisitions. these off-springs were further subdivided into 4 groups of 6 animals. The name of each group describes first, the condition of the fathers and second, the condition of their off-springs: Control-Control (CC). Control-Stress (CS). Stress-Control (SC) and SS (Stress-Stress). The long-term effects of peripubertal stress were examined when animals were 3 to 4 months-old.

Rats (n=6 per group. 495 ± 56 g) were anesthetized using isoflurane (2-3% in O_2_) during MRI and ^1^H-fMRS. Each rat was transported from the animal facility to the animal preparation room individually. Care was taken to manipulate each animal with very quiet movements. Each rat was tail vein cannulated in order to receive 2-Hydroxypropyl-β-cyclodextrine-cyclodextrine (HBC) followed by Corticosterone (CORT) at 2.5mg/kg (Sigma-Aldrich. Switzerland) in order to imitate the dose of steroid hormones in the plasma induced by substantial stress. Injection of HBC was performed 40 minutes before CORT in order to permeabilize the blood brain barrier to CORT. Each rat was placed in a dedicated stereotactic rat holder under continuous isoflurane anesthesia (2% in O_2_). The rat body temperature was monitored continuously by a rectal probe and maintained under physiological temperature (37±1°C) using warm circulating water around the animal. After each MR session. each animal was gently put back in its cage. When completely awake. the rat was returned to the animal facility.

### ^1^H-functional MRS in septal areas of the rat brain

All the experiments were performed on an actively shielded 9.4T/31cm bore magnet (Magnex. Varian) with 12cm gradients (400mT/m in 120μs). A quadrature Transmit/Receive 17mm surface coil was used. First and second order shims were adjusted using FASTMAP (20) in a 3×3×3 mm^3^ volume of interest placed over the lateral septum by reference to the Paxinos atlas (21) with a 3D-Gradient Echo sequence. In the voxel of interest (VOI). the water peak acquisitions resulted in water linewidths of 15 ± 3 Hz. Localized proton MR spectroscopy was performed using SPin ECho, full Intensity Acquired Localized (SPECIAL) MR spectroscopy (22). 40 blocks of 16 FIDs were acquired during basal conditions (as reported in ref (7)) followed by a bolus of HBC and an acquisition of 40 blocks of 16 FIDs and a bolus of CORT and another acquisition of 40 blocks of 16 FIDS equivalent to 40 minutes of acquisition. For absolute quantification. water signal was acquired using identical parameters without water suppression and using an average of 8 scans.

### Data analysis

The 16 × 40 FIDs were individually Fourier transformed. realigned and rephased and finally averaged for each animal for the basal, HBC and CORT conditions. The *in-vivo* ^1^H MR spectra were processed using LCModel (23). Absolute metabolite concentrations were obtained using unsuppressed water signal as a reference. In this study. 18 metabolites concentrations were quantified using databases of simulated spectra of metabolites (22) and measured macromolecular baseline. Averaged absolute concentration neurochemical profiles (NPs) were obtained for each group of rats. Finally. analysis of NPs was reduced to the main neurotransmitters (Glu, GABA) and metabolites (Gln, Glc, Lac) as well as neuronal marker (NAA) for an easier exploration. Changes in NPs were obtained by calculating 1) the relative change in absolute concentrations of Glu, Gln, Glc, Lac and NAA with regards to the basal concentrations of these metabolites and 2) the relative change in absolute concentrations of Glu, Gln, Glc, Lac and NAA with regards to post-CORT concentrations of these metabolites in the CC group of rats.

For the comparison of the shape or pattern described by neurochemical profiles using a quantifiable metric, the neurochemical profiles were considered as distributions of metabolites and the similarity between distributions was evaluated using a chi-square distance algorithm (Pattern recognition) (24) using a Matlab routine (http://vision.ucsd.edu/~pdollar/toolbox/doc/index.html). A parameter d was defined: if d was close to 0. histograms were considered very similar and the more the value d increased. the more histogram shapes were considered di-similar.

Time courses of metabolites were obtained for each individual and averaged across groups of rats: at bolus injection. at 10 minutes (averaging 10 × 16 FIDs). at 20 minutes (averaging the next 10 × 16 FIDs). at 30 minutes (averaging the next10 × 16 FIDs) and finally at 40 minutes (averaging the next10 × 16 FIDs). Relative changes in % were obtained relative to basal levels and relative to post-CORT levels in the CC group of rats.

### Statistical Analysis

All data are presented as mean ± standard error of the mean (S.E.M) unless otherwise stated. The data were analyzed statistically using two-way ANOVA to compare metabolite concentrations between groups. Post hoc tests were performed with the Bonferroni test. A pvalue< 0.05 was considered significant. The statistical analysis was performed using GraphPad Prism (Version 5. GraphPad Software Inc. La Jolla. USA).

## RESULTS

### CORT changes the metabolism of PS animals

The effects of CORT injection in pPS and iPS rats were evaluated by comparing post-CORT metabolite concentrations to pre-CORT metabolite concentrations in the rat septal area (Table1). The neurochemical profiles (NP) of the septal area for CC, CS, SC and SS groups of rats were obtained from the LCModel fitting of MR spectra **(Fig.2)** (including the reliable absolute quantification of 12 metabolite concentrations (CRLB below 20%) pre-and post-CORT challenge. A two-way ANOVA test revealed a global effect of CORT on NPs of CC and SC groups of rats compared to basal levels (p<0.0001). There was also a significant effect of pPS and iPS on groups of CORT-injected rats (p<0.0001). Exogenous CORT levels increased Glc levels in all groups of rats (CC: +34%; CS: +42% (p<0.05); SS: +13%) except for pPS rats (SC: −18%). Lac levels were reduced with identical proportions due to CORT in CC, SC and SS groups but not in CS rats (+22%). CORT elevated GABA levels in CS (+ 87%) and SS (+43%) rats but not in CC and SC rats (−19 and −25% respectively).

**Table 1:** Median and Mean (±Standard Deviation) Concentrations of Glutamate (Glu), Glutamine (Gln), γ-amino-butyric acid (GABA), Glucose (Glc), Lactate(Lac) and N-acetyl-aspartate (NAA) in the Septum of control and stressed rats under basal and corticosterone condition.

| $\mu\text{mol/g}$ | BASAL | | | | | | | | Corticosterone | | | | | | | |
| --- | --- | --- | --- | --- | --- | --- | --- | --- | --- | --- | --- | --- | --- | --- | --- | --- |
|  | CC |  | CS |  | SC |  | SS |  | CC |  | CS |  | SC |  | SS |  |
| **Glu** | 7.35 | 7.58±0.9 | 8.39 | 8.34±0.6 | 7.35 | 7.36±0.6 | 8.52 | 8.58±0.9 | 7.54 | 7.9±0.9 | 7.69 | 7.5±0.9 | 5.77 | 6.4±0.9 | 8.27 | 7.92±1.0 |
| **Gln** | 3.76 | 3.66±0.2 | 3.92 | 4.07±0.7 | 3.61 | 3.55±0.9 | 3.7 | 3.55±0.9 | 3.28 | 3.4±0.6 | 3.32 | 3.3±0.9 | 4.03 | 4.0±0.6 | 3.79 | 3.99±0.4 |
| **GABA** | 1.32 | 1.48±0.3 | 0.62 | 0.75±0.6 | 1.65 | 1.59±0.5 | 0.89 | 0.98±0.6 | 1.02 | 1.2±0.4 | 1.34 | 1.4±0.3 | 1.2 | 1.2±0.5 | 1.17 | 1.40±0.3 |
| **Glc** | 2.4 | 2.62±1.0 | 2.05 | 3.10±0.9 | 3.56 | 3.92±0.9 | 3.36 | 2.80±0.9 | 2.68 | 3.5±1.3 | 4.9 | 4.4±1.8 | 3.47 | 3.2±0.6 | 2.33 | 3.15±1.1 |
| **Lac** | 1.32 | 1.22±0.3 | 1.2 | 1.64±0.7 | 1.47 | 1.60±0.6 | 2.17 | 1.84±0.7 | 0.94 | 1.1±0.4 | 2.05 | 2.0±0.4 | 1.07 | 1.3±0.5 | 1.56 | 1.59±0.2 |
| **NAA** | 7.77 | 7.44±0.9 | 8.29 | 8.65±0.8 | 7.7 | 7.47±0.7 | 8.29 | 8.30±0.5 | 7.57 | 7.7±0.5 | 8.62 | 8.4±0.5 | 6.94 | 7.1±0.4 | 7.94 | 8.39±0.7 |

**Figure 2:**
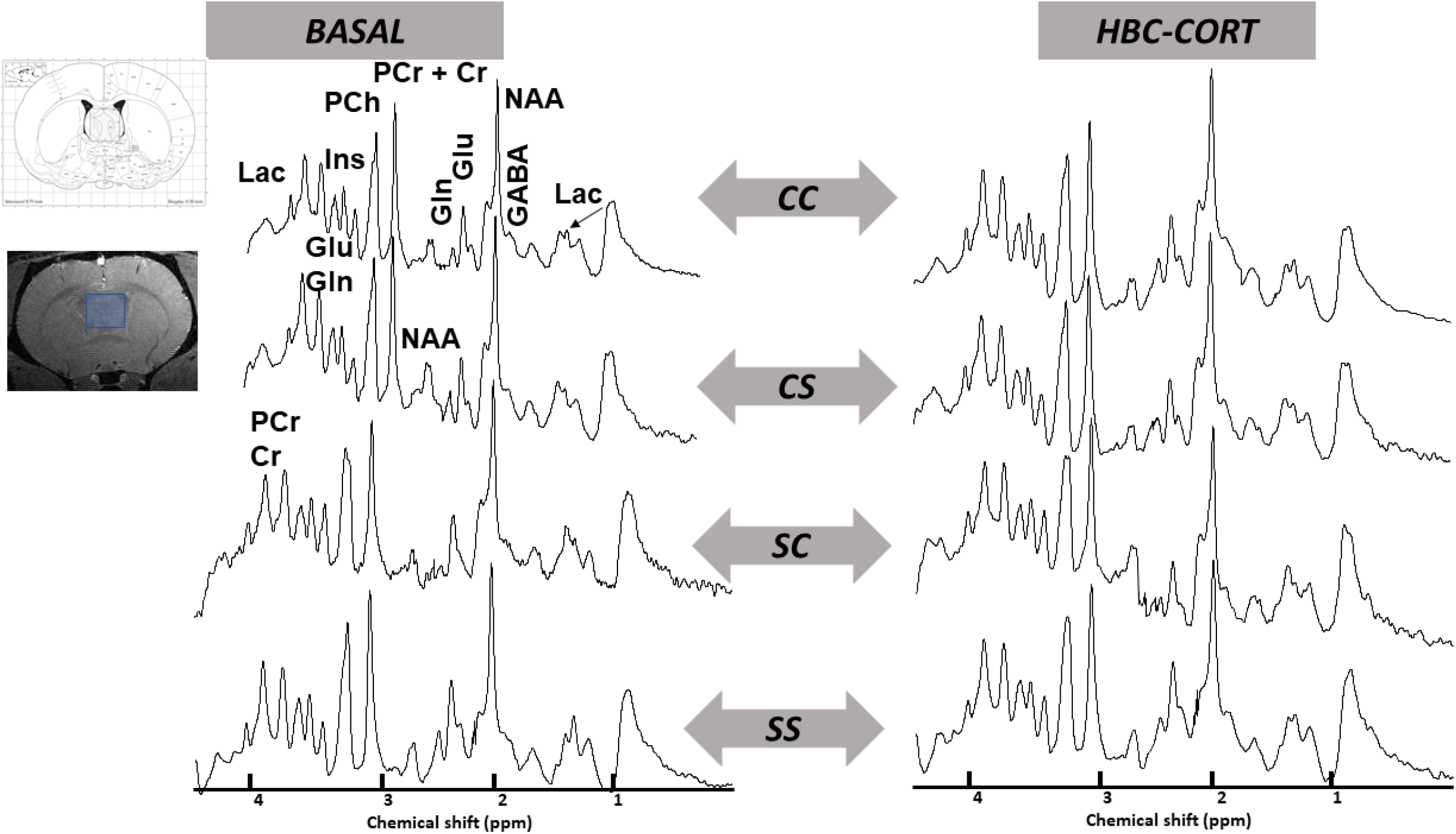
**Representative in-vivo ^1^H-MR spectra at 9.4T in the rat septal area during basal state and Corticosterone challenge localized on** an axial 3D T1-weighted gradient echo image of the rat brain and 27 μl voxel of interest in blue over the septal area delineated by reference to the Paxinos atlas (21) at a 0.3 mm distance from the bregma. Control-Control (CC) with labels. Control-Stress (CS). Stress-Control (SC) and Stress-Stress (SS). Glutamate (Glu). Glutamine (Gln). Lactate (Lac). N-Acetyl-Aspartate (NAA). Phosphocreatine (PCr). Creatine (Cr). Phosphocholine (PCho). Myo-inositol (Ins). Alanine (Ala). Glutathione (GSH). γ-aminobutyric acid (GABA). Taurine (Tau). Total Creatine (PCr + Cr).

### Classification of neurochemical profile changes in pPS and iPS rats

Important differences were observed visually on the NPs of each group due to CORT (**Fig.3**). NPs of the changes in metabolite concentrations due to CORT ((post-CORT-pre-CORT)/pre-CORT) for each group (**Fig. 3A**) were classified from the most similar to the least similar as follows: dcs-ss<dsc-ss<dcc-sc<dcc-ss<dcc-cs<dcs-sc: 46.68<53.78<167.94<206.08<213.21<741.09. NPs modified by CORT were the most different between CS and SC groups and the most similar between CS and SS. Changes relative to basal conditions were significant for all metabolites between CC and all other groups (p<0.001). Interestingly, only changes in Glu and NAA, which are both neuronal markers, were non-significant between CS and SC and between CS and SS.

**Figure 3:**
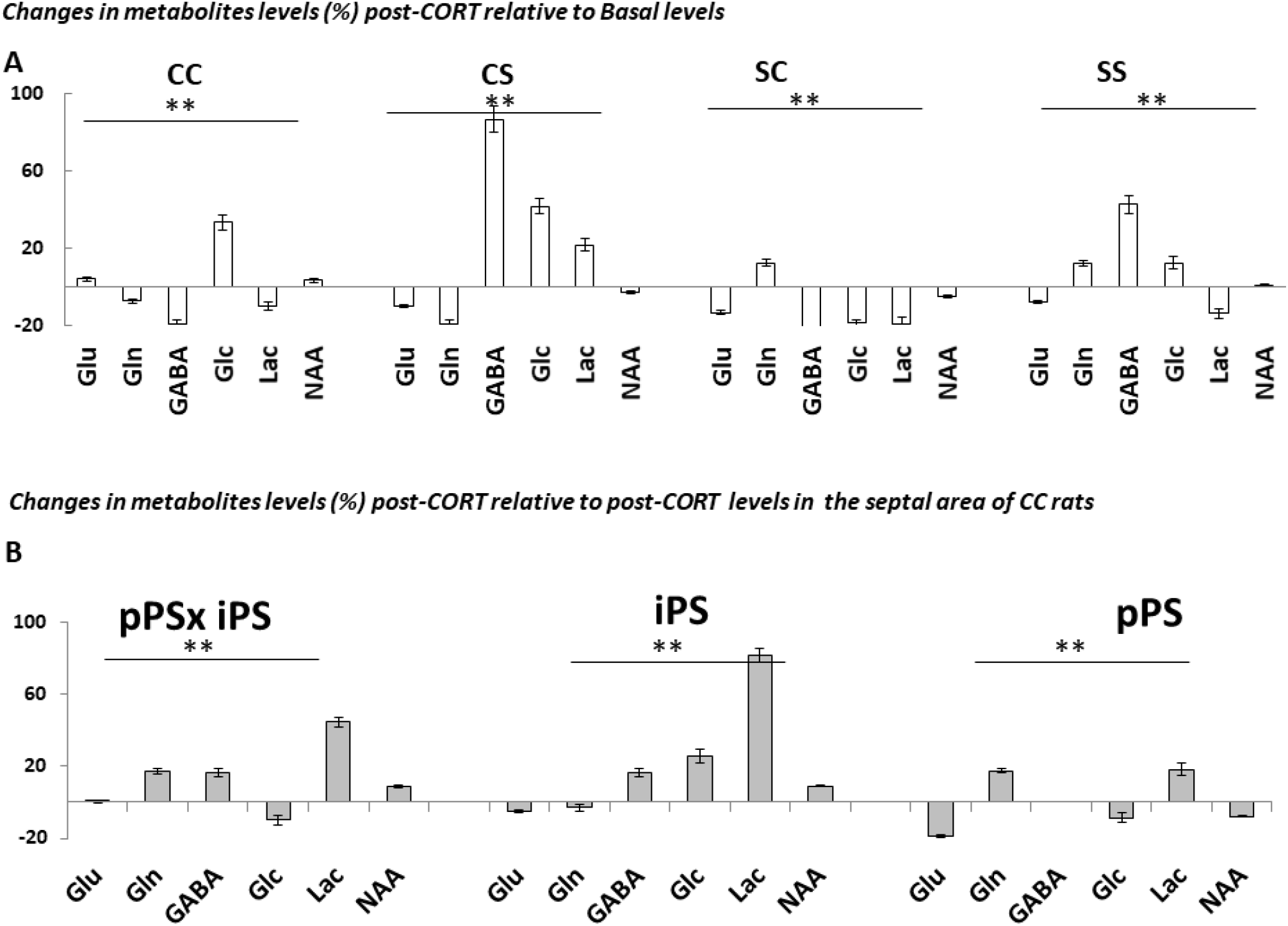
Neurochemical profile changes in metabolite concentrations. **A.** Changes (%) of metabolite concentrations due to CORT relative to basal metabolite concentrations for CC. CS. SC and SS groups. ** p< 0.001 denotes significant differences between groups. Concentration changes were not different between CS and SC for Glu (p>0.05) and between CS and SS for NAA (p>0.05). B. Changes (%) of metabolite concentrations due to CORT relative to post-CORT concentrations in CC. Neurochemical profile changes were compared to understand the influence of paternal PS only (SC) and individual PS (CS) only and their interaction (iPS × pPS = SS) on metabolite concentrations following CORT injection. ** p< 0.001 denotes significant differences between groups

The same strategy was applied for the comparison between post-CORT groups and post-CORTCC (**Fig 3B**): dpostCC-postCC< dpostCS-postSS< dpostSC-postSS< dpostSC-postCS (0<115.2<247.7<380.2). Differences in post-CORT neurochemical profiles of stress groups relative to the post-CORT control group (post-CORTCC) were again most pronounced between CS and SC groups demonstrating a differential action of pPS and iPS on corticosterone metabolism (**Fig. 3B**). A 2-way ANOVA statistical test with Bonferroni adjustment confirmed significant differences between all individual metabolite concentration changes relative to post-CORT CC metabolite concentrations of pPS and iPS groups of rats (p<0.001). Changes in Glu, Gln, Glc and Lac concentrations were also significant between rats affected by the interaction of pPS and iPS (pPS x iPS and rats affected by iPS only (p<0.001). Changes in Glu, GABA, Lac and NAA concentrations were also significant between rats affected by the interaction of pPS and iPS (pPS x iPS) and rats affected by pPS only (p<0.001). The comparison of stressed groups of rats post-CORT relative to the post-CORT CC group allowed us to evaluate the specific effects of pPS, iPS and their interaction during CORT challenge. Positive Lac concentration changes were found in all groups. These changes were more elevated in the CS (+82%) and SS (+45%) groups than in SC (+18%) group. On the other hand, Glc levels decreased by 10% in SC and SS groups. GABA levels remained unchanged in SC rats. GABA levels increased by 17% together with increased Glc levels by 26% in CS rats.

### CORT-induced metabolic concentration changes in PS rats as a function of time

Changes in concentrations of Glu, Lac, Glc, NAA as well as PCr/ Cr were obtained as a function of time. CORT had effects on both septal neuro-energetics and neurotransmission.

#### Qualitative approach

**Fig 4** shows the evolution of Glc and Lac concentrations for one typical rat of each group. 10 minutes post-CORT injection, Glc concentrations were increased in CC and CS rats while levels of Glc were already elevated in SC and SS rats. Lac levels also presented higher values in these rats. The Glc to Lac ratio was calculated for each rat. CORT injection induced a significant increase of the Glc to Lac ratio at 10 minutes in CC and 20 minutes in SS. The Glc to Lac ratio remained constant up to 40 minutes after CORT injection in CC and SS, while the ratio remained low in CS and SC rats without significant difference between the two animals. Although qualitative, these observations suggest adaption of metabolism in septal areas in SS rats. In addition, the PCr to Cr ratio was lower in CS, SC and SS rats compared to values in the CC rat suggesting an enhanced use of energy in PS rats compared to control rats. Finally, the Glu to Gln ratio was calculated showing lower levels in CS, SC and SS rats compared to levels in the CC rat after HBC and CORT injections thus suggesting alterations of neurotransmission due to CORT in PS rats.

**Figure 4.**
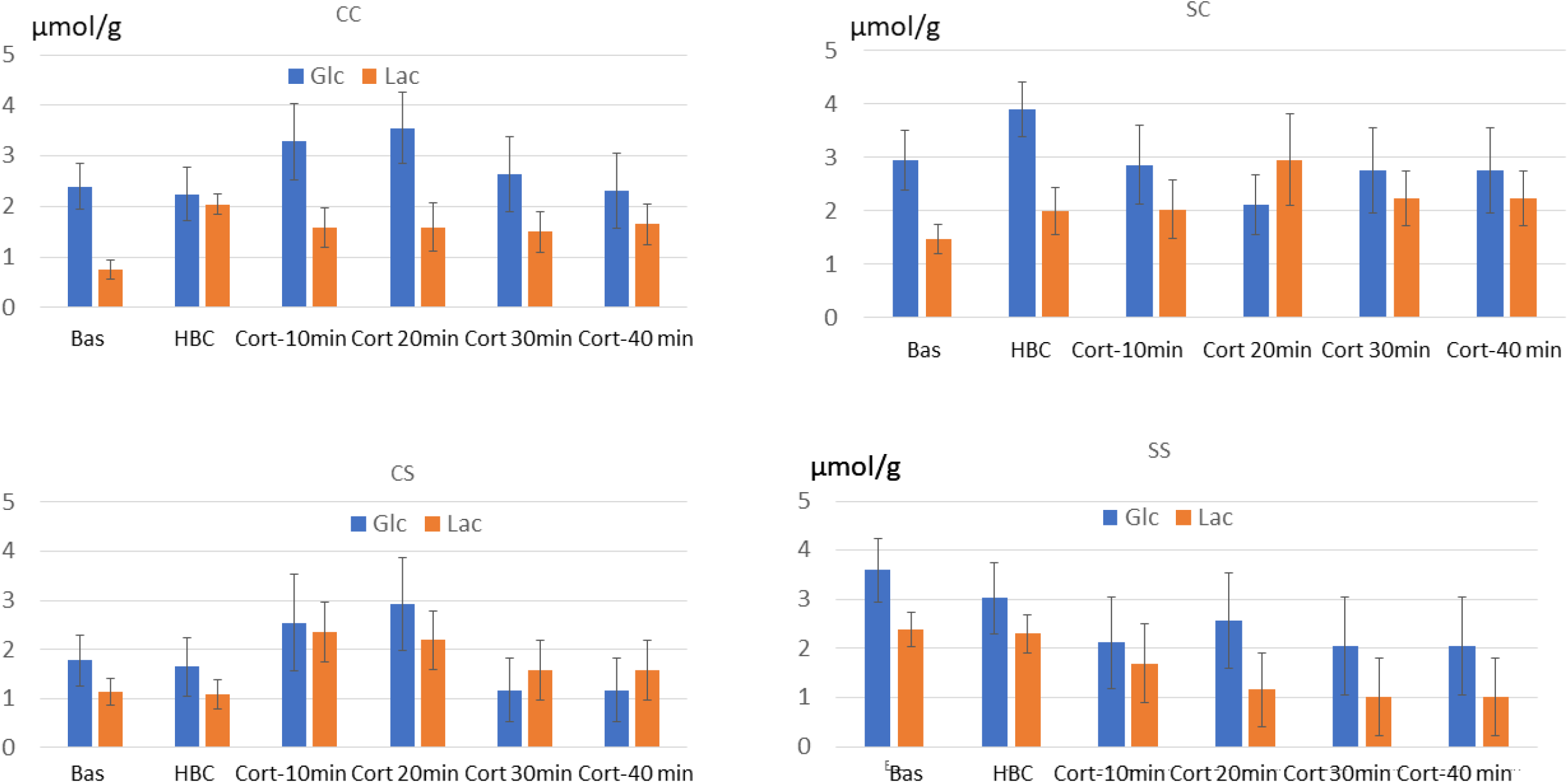
Qualitative analysis: Typical evolution of Glc and Lac concentrations (μmol/g ± CRLB) for basal level, after HBC injection and after CORT injection at 10 min, 20min, 30 min and 40 min in one typical rat pertaining to each group.

#### Quantitative approach and comparison between groups of rats as a function of time

In order to identify the quantitative changes induced by CORT on septal metabolites in the different groups of rats, metabolite concentration changes were calculated relative to the basal levels. These changes are illustrated in **Fig.5**. Significant differences were found between groups for Glc, Lac, Glu and the Glc to Lac ratio (Table 2) at all time points.

**Figure 5.**
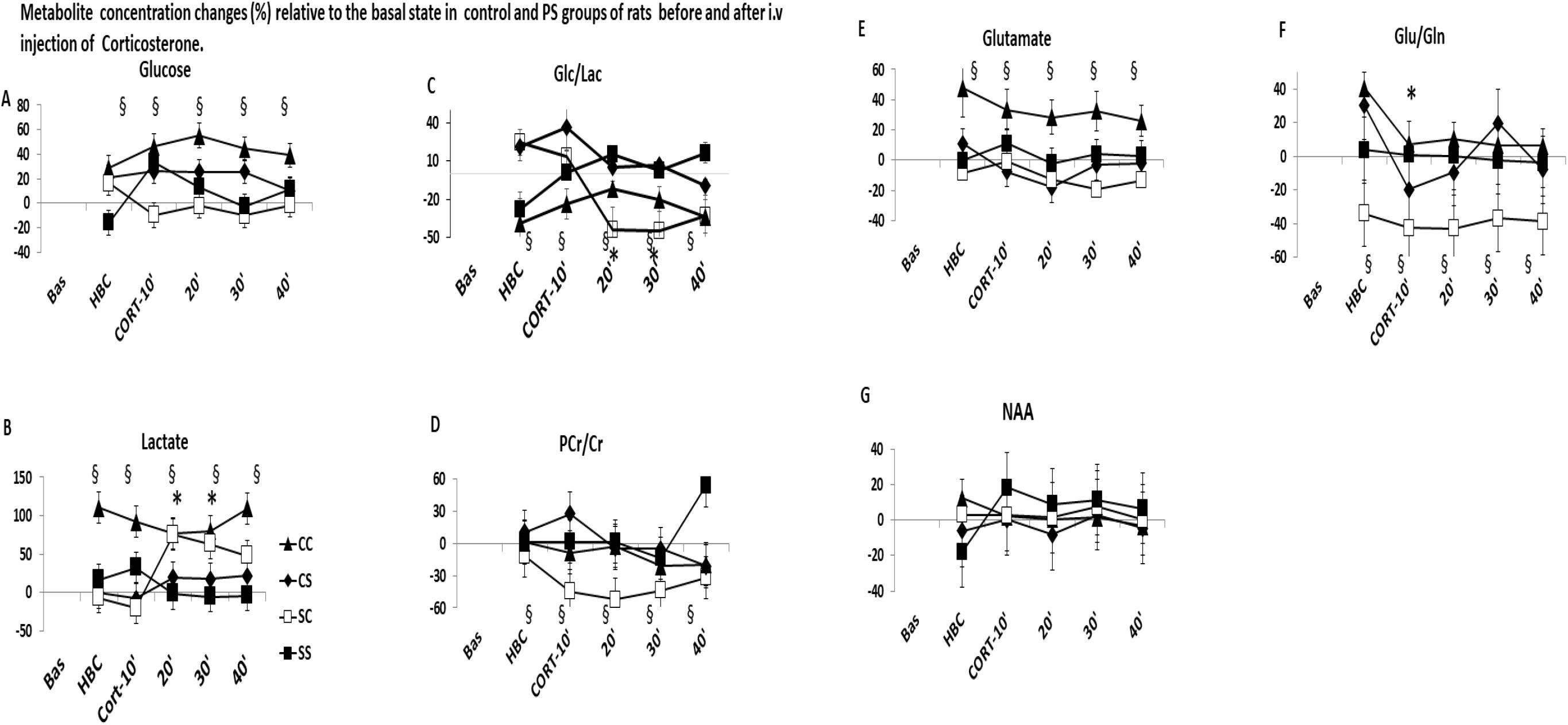
**Percent change of metabolite concentration as a function of time relative to metabolite concentrations at basal levels for each group of rats** for A: Glucose; B: Lactate; C: Glc/Lac; D: PCr/Cr; E: Glutamate; F: Glu/Gln; G: NAA. § indicates significant differences between groups as described in Table 2. Pvalues for the comparison between timecourses for each metabolite are presented in Table 2. * indicates significant change between timepoints.

**Table 2:**
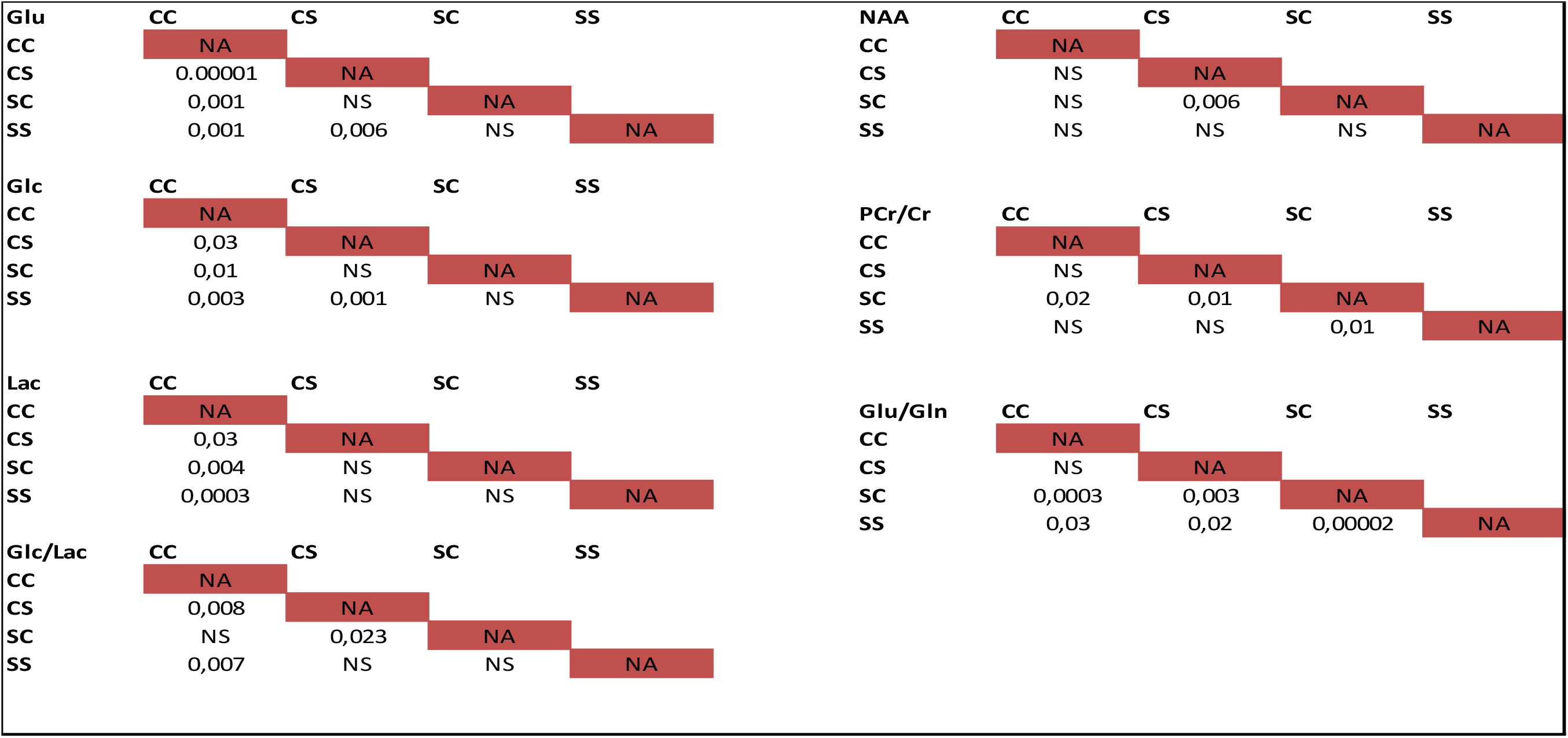
Statistics: Comparison of metabolite concentration changes between groups of rats relative to the basal state (No HBC, NO CORT) for Glutamate (Glu), Glucose (Glc), Lactate (Lactate), N-Acetyl-Aspartate (NAA), Gl to Lac ratio, Phosphocreatine to creatine (PCr/Cr) and glutamate to Glutamine ratio (Glu/Gln). Pvalues are reported.

| **Glu** | **CC** | **CS** | **SC** | **SS** | **NAA** | **CC** | **CS** | **SC** | **SS** |
| --- | --- | --- | --- | --- | --- | --- | --- | --- | --- |
| **CC** | NA |  |  |  | **CC** | NA |  |  |  |
| **CS** | 0,00001 | NA |  |  | **CS** | NS | NA |  |  |
| **SC** | 0,001 | NS | NA |  | **SC** | NS | 0,006 | NA |  |
| **SS** | 0,001 | 0,006 | NS | NA | **SS** | NS | NS | NS | NA |
| **Glc** | **CC** | **CS** | **SC** | **SS** | **PCr/Cr** | **CC** | **CS** | **SC** | **SS** |
| **CC** | NA |  |  |  | **CC** | NA |  |  |  |
| **CS** | 0,03 | NA |  |  | **CS** | NS | NA |  |  |
| **SC** | 0,01 | NS | NA |  | **SC** | 0,02 | 0,01 | NA |  |
| **SS** | 0,003 | 0,001 | NS | NA | **SS** | NS | NS | 0,01 | NA |
| **Lac** | **CC** | **CS** | **SC** | **SS** | **Glu/Gln** | **CC** | **CS** | **SC** | **SS** |
| **CC** | NA |  |  |  | **CC** | NA |  |  |  |
| **CS** | 0,03 | NA |  |  | **CS** | NS | NA |  |  |
| **SC** | 0,004 | NS | NA |  | **SC** | 0,0003 | 0,003 | NA |  |
| **SS** | 0,0003 | NS | NS | NA | **SS** | 0,03 | 0,02 | 0,00002 | NA |
| **Glc/Lac** | **CC** | **CS** | **SC** | **SS** |  |  |  |  |  |
| **CC** | NA |  |  |  |  |  |  |  |  |
| **CS** | 0,008 | NA |  |  |  |  |  |  |  |
| **SC** | NS | 0,023 | NA |  |  |  |  |  |  |
| **SS** | 0,007 | NS | NS | NA |  |  |  |  |  |

### Glucose changes

Glc levels were increased in CC and CS groups and remained constant throughout the entire HBC-CORT challenge while Glc level changes increased significantly at 10 min (p < 0.01) post-CORT injection before decreasing again in SS rats. In SC rats, Glc changes remained low during CORT challenge.

### Lactate changes

Relative Lac concentration changes were significant and constant during the entire HBC-CORT challenge in CC rats whereas significant changes occurred at 20 min post-CORT injection in SC rats. Changes remained lower for CS and SS groups for which no changes relative to basal levels were detected from 20 min post-CORT.

### Glc to Lac index

The Glc to Lac ratio remained the lowest throughout the entire HBC-CORT challenge for CC rats with decreased levels compared to basal levels at all timepoints. In SS rats, the Glc to Lac index followed the same evolution becoming positive at 20 min post-CORT and then falling down again. In CS and SC rats, the Glc to Lac index was increased post HBC and at 10 min post CORT before decreasing significantly for both groups at 20 min post-CORT. This index became negative and constant up to 40 min post-CORT. Interestingly, the Glc to Lac index demonstrated two different patterns of evolution during HBC-CORT challenge: similar time-course shapes for CS and SC on the one hand and for CC and SS on the second hand.

### Glu, NAA and Glu/Gln ratio

Concentration changes of Glu and NAA and allowed the evaluation of CORT effects on neurotransmission since both metabolites are neuronal markers. The Glu to Gln ratio as a function of time was also evaluated. Interestingly, concentration changes in CC as a function of time were significantly different from all the other groups and no significant difference was found between CS, SC and SS rats. In CC group, mean changes relative to basal levels remained constant (+ 35%) all along the HBC-CORT challenge. By contrast, no significant differences were found between all groups regarding NAA changes, which remained nearly unchanged compared to basal levels for all CC, Cs and SC groups. A 20% increase relative to basal levels was noticed in CC 10 minutes after CORT injection. In this group NAA changes remained positive until the end of the challenge.

Relative to basal levels the Glu to Gln ratio decreased significantly in SC rats and remained lower compared to all the other groups during the entire HBC-CORT challenge. There were no changes in Glu to Gln in SS rats during the entire HBC-CORT challenge while changes remained also low in CC and CS groups after a significant increase at HBC injection. Changes were significantly different between all groups except between CC and CS.

The specific effects of pPS and iPS on metabolite dynamics during CORT challenge were assessed relative to the control group. Metabolite concentrations in CS, SC and SS groups were normalized to metabolite concentrations in CC (**Fig. 6**). No significant changes were found between groups for Lac and Glu (p>0.05). However, both Glu and Lac relative concentrations decreased significantly (p<0.01) after HBC injection remaining low and constant during the entire CORT challenge for Glu. In SS rats, Lac levels remained unchanged compared to those in the CC group while Lac levels increased again 20 minutes after CORT injection in CS and SC groups. Glc levels were higher in SC and CS rats and remained unchanged in SS rats compared to CC rats at the basal state. During HBC-CORT challenge, relative Glc levels were decreased in all three groups. Glc levels remained higher in CS than in CC rats (from +30 % at basal state to +18 % 10 min post-CORT and + 2 % 40 min post-CORT). On the contrary, significant changes occurred after 10 min post-CORT in SC and after HBC injection in SS (p<0.001) for which lower Glc levels were detected compared to CC rats. The relative change in Glc to Lac index demonstrated significant differences after HBC injection and 10-min post-CORT injection showing a highest index for SC and a lowest index for SS. No differences between groups were noticed from 20-min post-CORT to the end of the challenge.

**Figure 6.**
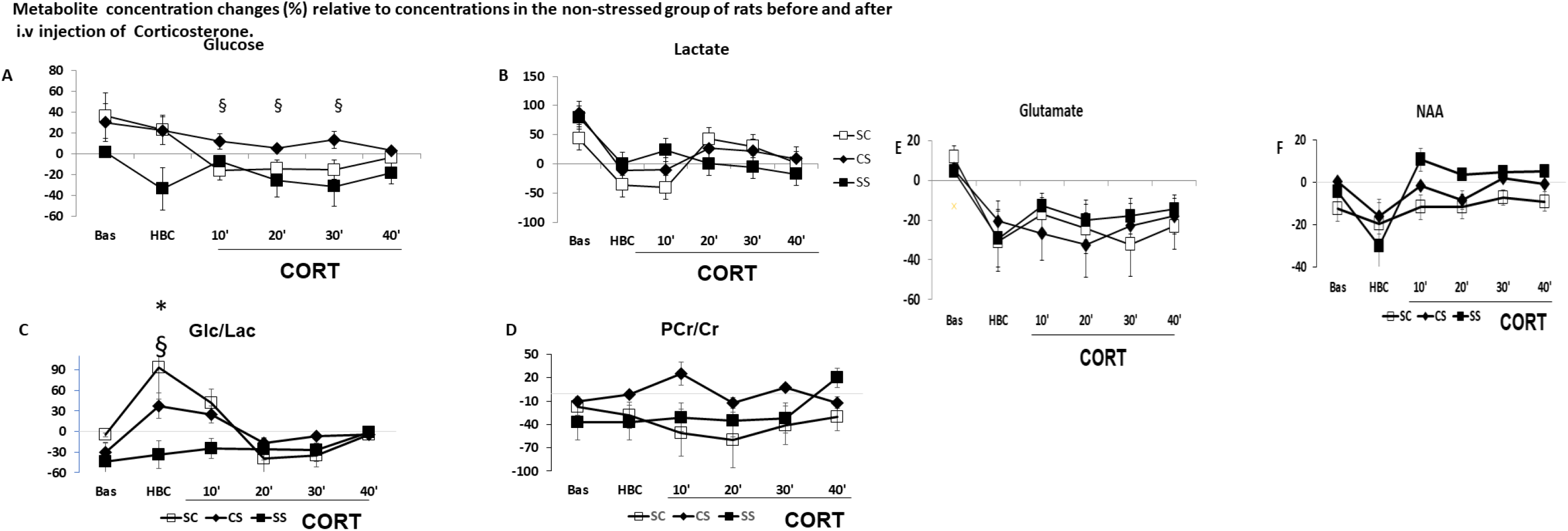
Percent change of metabolite concentration post-CORT relative to post-CORT CC metabolite concentrations. § indicates significant difference between groups and * indicates significant differences between timepoints (§ p< 0.05; * p< 0.05).

The PCr/Cr ratio was significantly decreased in SC and SS rats relative to CC rats at all time points but remained unchanged on average in CS rats. HBC injection created a significant depression of NAA levels relative in all groups relative to CC rats. There was a significant difference between SC and SS relative NAA levels after CORT injection: in SS rats NAA changes remained positive until the end of the CORT challenge while NAA changes returned to levels of change before HBC injection and remained negative in SC rats. CS rats also demonstrated negative NAA changes during basal stages and CORT challenge.

## DISCUSSION

A better knowledge of the underlying mechanisms of stress disorders is necessary for a better understanding of the evolution of these disorders into psychiatric diseases. In this context, an enhanced characterization of the underlying brain metabolism may be of great interest. Here, the changes in energy metabolism and neurotransmission of the septal area of PS rats post-corticosterone challenge using dynamic ^1^H-MRS were characterized. In the context of the present study, the term “^1^H-fMRS” was preferred since ELS and the subsequent hormonal alterations are known to induce changes in neuronal activity in multiple areas of the brain linked to changes in metabolism (25, 26). Here, ^1^H-fMRS was conducted assuming that PS and exogenous CORT can be considered as endogenous stimulators of septal metabolism in rats.

fMRS is a powerful technique enabling the quantitative estimation of metabolic changes between periods of brain activity and periods of rest. ^1^H-fMRS demonstrated enormous potential for investigating the human and rodent brain neuro-energetics upon external stimulation (13, 27, 28, 29) and is becoming a useful complementary technique to neuroimaging techniques (fMRI, PET) for the *in-vivo* exploration of the neurochemical consequences of cognitive tasks (30). ^1^H-fMRS is gradually becoming a technique of choice for exploring functional neurochemical changes in neuro-pathologies (15, 16). This technique can measure the dynamics of important neurotransmitters *in-vivo* such as Glu and GABA and is therefore a technique of interest for studies on pharmacological challenge and neurobiological mechanisms (30).

The present study confirmed that PS induces significant metabolic alterations in the septal areas of the rat (7). Moreover, CORT induced differential effects on septal metabolism whether it was delivered to Control or PS animals and investigations using fMRS helped discriminating between individually-stressed and paternally stressed animals. Importantly, metabolic adaption to CORT was assessed. This study also revealed the importance of considering multiple metabolite changes simultaneously in order to address specific differences between individually and paternally-stressed rats as well as interactions between the two. Therefore, neurochemical profiles were classified using a dissimilarity metric and temporal neurochemical profile changes were compared between groups. Classification methodologies and pattern recognition techniques have already demonstrated their potential for the analysis of spectroscopic data (31). Nevertheless, few studies have been conducted outside of the field of oncology, essentially because of the need to define specific criteria for an adequate characterization of neurochemical profiles including ways to discard low signal to noise ratio or inadequately shimmed spectra (32).

Qualitatively, we examined the similarity of neurochemical profiles between groups of rats using a metric of distance used in pattern recognition algorithms. This method determined that neurochemical profile changes were very di-similar upon CORT challenge between iPS (CS) and pPS (SC) rats. However, counter-intuitively, neurochemical profile changes were more similar between CS and SS and between SC and SS, which were confirmed upon temporal analysis of neurochemical changes. It is often difficult to compare up to 22 simultaneously acquired and quantified metabolites. Moreover, concentration changes in one metabolite are often linked to changes in another such as for example Glu, Gln and GABA. Here, we proposed to consider the neurochemical profiles as distributions of metabolites in order to explore the overall effects of CORT on septal metabolism. While this approach remains only qualitative and needs to be tested further and validated, we found it useful in the present study to extract overall effects of ELS on the metabolism of iPS and pPS rats.

Glc levels were increased in all groups of rats relative to basal levels following HBC-CORT injections. It is known that individuals repeatedly exposed to glucocorticoids present high blood Glc levels including in situations with no stress and a prevalent disposition to diabetes (10).In individual SC and SS rats, elevated basal Glc levels were noticed compared to CC and CS rats. This could result from already higher CORT levels in the descendants of peripubertally stressed animals.

Due to stress, increased levels of energy are necessary suggesting that other sources of energy than Glc are necessary. In this context, several studies demonstrated increased levels of Lac due to stress and adaptions qualified as “cerebral demands in Lac”. In addition, other studies mentioned anti-depression effects of Lac, which could act as a hormone improving the effectiveness of metabolism during stress.

Relative to basal levels, Lac concentrations increased in all groups. However, the highest increases were seen for the CC group while only mild increases were seen at 10 min post-CORT for SS rats. Relative to CC, Lac concentrations were elevated in CS, SC and SS groups during basal stages. Lac concentration changes were reduced in all groups relative to CC upon HBC and CORT injections but remained almost unchanged in SS rats only. We hypothesize that relative Glc and Lac concentration changes suggest a differential neuro-energetic modulation of septal areas between PS groups, which could depend on the equilibrium between Glc and Lac concentrations. A Glc to Lac index was calculated, allowing to recognize similarities between groups of rats.

Stress, whether acute or chronic, demonstrated differential effects on neurotransmission (33). Acute secretions of CORT induce increases in neuronal receptor mobility, neuronal plasticity and allow faster adaption to cerebral activity demands. However, when stress becomes chronic and CORT secretions are prolonged, neurotransmission is significantly altered. Notably, previous studies in rodents demonstrated reduced NAA, Glu, Gln and GABA concentrations (34). In the present study, NAA changes relative to the control group CC in CS and SC groups suggest alterations of neuronal function in septal areas. Interestingly, CORT seems to improve neuronal functions in SS. However, CORT altered glutamatergic neurotransmission in all PS affected rats relative to CC. In addition, the Glu to Gln index was unchanged for SS and significantly decreased for CS and SC relative to basal levels indicating reduced neurotransmission in these groups and adaption in SS.

### Pitfalls of the study

Previous studies revealed important interactions between glucocorticoids and the GABAergic system with important anxiolytic effects of GABAergic transmission (35, 36). In the present work, despite a decrease of GABA levels in all groups upon CORT injection in accordance with previously reported results (35), GABA quantification was not reliable (CRLBs > 40%). The investigation of other metabolites of interest such as: ascorbate, aspartate, glycine, phosphocholine, glutathione could help confirming results (34). However, increased sensitivity would be necessary. In addition, Gapp et al. (34) found important regional differences in response to ELS. These differences could also be present within nuclei of the same structure. A wide number of studies focused on the lateral septum response to ELS in rodents (36). This limbic structure plays an important role in anxiety and represents a relay between the cortex and the hypothalamic-pituitary-adrenal axis (HPA), which is significantly implicated in stress issues. Here, the voxel of interest encompasses all the different nuclei of the septal area and doesn’t allow specificity of results.

Increased Lac concentrations in the different groups of rats could also be attributed to prolonged MRS sessions under isoflurane anaesthesia (37). In the present case, this hypothesis cannot be excluded but CORT effects and PS effects were compared in 4 groups of 6 rats anesthetized under the exact same conditions and during the same duration. Moreover, relative concentrations were calculated minimizing the impact of anaesthesia on results.

A non-negligible impact of HBC was noticed on profiles as a function of time was observed. Cyclodextrins are used for the conception of treatments due to their hydrophilic properties enabling the solubility of many therapeutic compounds. In the present study, 2-hydroxypropyl-β-cyclodextrin was used to permeabilize the blood brain barrier to CORT. However, recent studies demonstrated that HBC can also be used as active molecules for the treatment of neurologic disorders (38, 39). Rats were manipulated with care during experiments but potential rise of CORT levels due to catheterization of caudal veins as well as injections of CORT and HBC are not excluded.

Finally, blood oxygen level dependent fMRI prior and post CORT would have allowed a better assessment of functional changes in septal areas and/or the assessment of changes in functional connectivity, which correlations with metabolic findings could be very informative while a better positioning of VOIs (39) could add more specificity to metabolite measurements.

### Perspectives

fMRS (or dynamic MRS) represents a technique of choice to investigate mechanisms associated to stress. The Glc to Lac index measured with ^1^H-fMRS could be an effective marker of PS response in rodents but remains to be validated using neuroimaging and immunohistochemical techniques. In addition, the investigation of the mechanisms underlying PS and their evolution into adulthood could have important benefits for understanding the development of schizophrenia as well as diabetes.

^1^H-fMRS was used in conjunction with external stimuli in order to examine metabolic responses related to cerebral activity in rodents (13, 40). Here, PS and Cort injection were also considered as stimuli modulating cerebral activity although not delivered in a controlled and periodic manner as electrical stimuli. Functional activity of corticosteroids and their receptors was already reported. ^1^H-fMRS can therefore be used to study long-term metabolic effects of stress demonstrating the enormous translational potential of this technique.

## Acknowledgements

The MR acquisitions of this study were conducted at the Centre d’Imagerie Médicale (CIBM) of UNIL. EPFL. HUG. CHUV and Leenards and Jeantet foundations.

## Notes

### Competing Interest Statement

The authors have declared no competing interest.

## References

[1] Summers CH. Yaeger JDW. Staton CD. Arendt DH. Summers TR. Orexin/hypocretin receptor modulation of anxiolytic and antidepressive responses during social stress and decision-making: Potential for therapy. Brain Res. 2018. pii: S0006-8993(18)30662-0. doi: 10.1016/j.brainres.2018.12.036

[2] Ulrich-Lai YM. Fulton S. Wilson M. Petrovich G. Rinaman L. Stress exposure. food intake and emotional state. Stress. 2015;18(4):381–99. doi: 10.3109/10253890.2015.1062981.

[3] Fonzo GA. Diminished positive affect and traumatic stress: A biobehavioral review and commentary on trauma affective neuroscience. Neurobiol Stress. 2018;9:214–230. doi:10.1016/j.ynstr.2018.10.002.

[4] Howes OD. McCutcheon R. Owen MJ. Murray RM.The Role of Genes. Stress. and Dopamine in the Development of Schizophrenia.Biol Psychiatry. 2017 Jan 1;81(1):9–20. doi: 10.1016/j.biopsych.2016.07.014.

[5] Ahn SN. Lee J. Effects of sensory awareness. imagery and observation on electroencephalography in adult with psychological stress.J Phys Ther Sci. 2019; 31(1):17–19. doi: 10.1589/jpts.31.17.

[6] Di Iorio CR. Carey CE. Michalski LJ. Corral-Frias NS. Conley ED. Hariri AR. Bogdan R. Hypothalamic-pituitary-adrenal axis genetic variation and early stress moderates amygdala function. Psychoneuroendocrinology. 2017;80:170–178. doi: 10.1016/j.psyneuen.2017.03.016.

[7] Cordero MI. Just N. Poirier GL. Sandi C. Effects of paternal and peripubertal stress on aggression. anxiety. and metabolic alterations in the lateral septum. European Neuropsychopharmacology 2016. 26 (2). 357–36

[8] Strasser A. Xin L. Gruetter R. Sandi C. Nucleus accumbens neurochemistry in human anxiety: A 7 T ^1^H-MRS study.Eur Neuropsychopharmacol. 2018. pii: S0924-977X(18)32006-6. doi: 10.1016/j.euroneuro.2018.12.015.

[9] Nemeroff CB. Neurobiological consequences of childhood trauma. J Clin Psychiatry. 2004;65 Suppl 1:18–28.

[10] Picard M. McEwen BS. Epel ES. Sandi C. An energetic view of stress: Focus on mitochondria. Front Neuroendocrinol. 2018;49:72–85. doi: 10.1016/j.yfrne.2018.01.001.

[11] Just N. Gruetter R. Detection of neuronal activity and metabolism in a model of dehydration-induced anorexia in rats at 14.1 T using manganese-enhanced MRI and ^1^H MRS.NMR Biomed. 2011;24(10):1326–36.

[12] Just N. Cudalbu C. Lei H. Gruetter R. Effect of manganese chloride on the neurochemical profile of the rat hypothalamus. J Cereb Blood Flow Metab. 2011;31(12):2324–33.

[13] Just N. Xin L. Frenkel H. Gruetter R. (2013).Characterization of sustained BOLD activation in the rat barrel cortex and neurochemical consequences.Neuroimage.;74:343–51. doi: 10.1016/j.neuroimage.2013.02.042.

[14] Schaller B. Mekle R. Xin L. Kunz N. Gruetter R. (2013). Net increase of lactate and glutamate concentration in activated human visual cortex detected with magnetic resonance spectroscopy at 7 tesla. J Neurosci Res. 91(8):1076–1083. http://dx.doi.org/10.1002/jnr.23194.

[15] Gussew A. Rzanny R . Erdtel M . Scholle HC. Kaiser WA . Mentzel H J. . Reichenbach JR. Time-resolved functional 1 H MR spectroscopic detection of glutamate concentration changes in the brain during acute heat pain stimulation NeuroImage 49 (2010) 1895–1902

[16] Mullins PG. Towards a theory of functional magnetic resonance spectroscopy (fMRS): A meta-analysis and discussion of using MRS to measure changes in neurotransmitters in real time. Scandinavian Journal of Psychology. 2018. 59. 91–103

[17] Joëls M,Sarabdjitsingh RA, Karst H 2012 Unraveling the time domains of corticosteroid hormone influences on brain activity: rapid. slow. and chronic modes. Pharmacological Reviews 64 901–938.

[18] Toledo-Rodriguez M. Sandi C. Stress during Adolescence Increases Novelty Seeking and Risk-Taking Behavior in Male and Female Rats.Front Behav Neurosci. 2011 Apr 7;5:17.

[19] Márquez C. Poirier GL. Cordero MI. Larsen MH. Groner A. Marquis J. Magistretti PJ. Trono D. Sandi C. Peripuberty stress leads to abnormal aggression. altered amygdala and orbitofrontal reactivity and increased prefrontal MAOA gene expression. Transl Psychiatry. 2013;3:e216. doi: 10.1038/tp.2012.144

[20] Gruetter R. Tkác I. 2000. Field mapping without reference scan using asymmetric echo-planar techniques. Magn. Reson. Med. 2000; 43:319–323.

[21] Paxinos G and Watson C. (2006). The Rat Brain in Stereotaxic Coordinates Sixth Edition. Academic Press. san Diego.

[22] Mlynárik V. Gambarota G. Frenkel H. Gruetter R.Localized short-echo-time proton MR spectroscopy with full signal-intensity acquisition. Magn Reson Med. 2006;56(5):965–70

[23] Provencher SW. Estimation of metabolite concentrations from localized in vivo proton NMR spectra. Magn Reson Med. 1993 Dec;30(6):672–9.

[24] Ying S. Wen Z. Shi J. Peng Y. Peng J. Qiao H. Manifold Preserving: An Intrinsic Approach for Semisupervised Distance Metric Learning. IEEE Trans Neural Netw Learn Syst. 2018;29(7):2731–2742. doi: 10.1109/TNNLS.2017.2691005. Epub 2017 May 18.

[25] Tzanoulinou S, Riccio O, de Boer MW, Sandi C. Peripubertal stress-induced behavioral changes are associated with altered expression of genes involved in excitation and inhibition in the amygdala. Transl Psychiatry. 2014 Jul 8;4(7):e410. doi: 10.1038/tp.2014.54.

[26] Papilloud A, Guillot de Suduiraut I, Zanoletti O, Grosse J, Sandi C. Peripubertal stress increases play fighting at adolescence and modulates nucleus accumbens CB1 receptor expression and mitochondrial function in the amygdala. Transl Psychiatry. 2018 Aug 15;8(1):156. doi: 10.1038/s41398-018-0215-6.

[27] Schaller B. Xin L. O’Brien K. Magill AW. Gruetter R (2014) Are glutamate and lactate increases ubiquitous to physiological activation? A (1)H functional MR spectroscopy study during motor activation in human brain at 7Tesla. Neuroimage 93(Pt 1):138–145. http://dx.doi.org/10.1016/j.neuroimage.2014.02.016

[28] Sonnay S. Poirot J. Just N. Clerc AC. Gruetter R. Rainer G. Duarte JMN. (2018). Astrocytic and neuronal oxidative metabolism are coupled to the rate of glutamate-glutamine cycle in the tree shrew visual cortex. Glia. ;66(3):477–491. doi: 10.1002/glia.23259.

[29] Sonnay S. Duarte JMN. Just N. (2017). Lactate and glutamate dynamics during prolonged stimulation of the rat barrel cortex suggest adaptation of cerebral glucose and oxygen metabolism. Neuroscience. 2017a;346:337–348. doi:10.1016/j.neuroscience..01.034.

[30] Woodcock EA. Anand C. Khatib D. Diwadkar VA. Stanley JA. Working Memory Modulates Glutamate Levels in the Dorsolateral Prefrontal Cortex during 1 H fMRS. Front. Psychiatry. 2018. doi: 10.3389/fpsyt.2018.00066

[31] Zarinabad N, Abernethy LJ, Avula S, Davies NP, Rodriguez Gutierrez D, Jaspan T, MacPherson L, Mitra D, Rose HEL, Wilson M, Morgan PS, Bailey S, Pizer B, Arvanitis TN, Grundy RG, Auer DP, Peet A. Application of pattern recognition techniques for classification of pediatric brain tumors by in vivo 3T (1) H-MR spectroscopy-A multi-center study. . Magn Reson Med. 2018 Apr;79(4):2359–2366. doi: 10.1002/mrm.26837.

[32] M Wilson, O Andronesi, PB Barker, R Bartha, A Bizzi, PJ Bolan,… Methodological consensus on clinical proton MRS of the brain: Review and recommendations Magnetic resonance in medicine. 2019 82 (2), 527–550

[33] Gill K M, Grace AA .Differential effects of acute and repeated stress on hippocampus and amygdala inputs to the nucleus accumbens shell. Int J Neuropsychopharmacol. 2013 Oct;16(9):2013–25. doi: 10.1017/S1461145713000618.

[34] Gapp k. Corcoba A. Van Steenwyk G. Mansuy IM. Duarte JMN. Brain metabolic alterations in mice subjected to postnatal traumatic stress and in their offspring. 2017. Journal of Cerebral Blood Flow & Metabolism 37 (7). 2423–2432

[35] Mody I. Maguire J.The reciprocal regulation of stress hormones and GABA(A) receptors.Front Cell Neurosci. 2012 Jan 30;6:4. doi: 10.3389/fncel.2012.00004

[36] Hodges TE, Louth EL, Bailey CDC, McCormick CM Adolescent Social Instability Stress Alters Markers of Synaptic Plasticity and Dendritic Structure in the Medial Amygdala and Lateral Septum in Male Rats., Brain Struct Funct, 2019 ; Vol. 224, pp. 643–659.

[37] Just N. Proton functional magnetic resonance spectroscopy in rodents. NMR in Biomedicine, 2020;e4254

[38] Vecsernyés M, Fenyvesi F, Bácskay I, Deli MA, Szente L, Fenyvesi É. Cyclodextrins, Blood-Brain Barrier, and Treatment of Neurological Diseases. Arch Med Res, 2014 ;Vol. 45, pp. 711–729.

[39] Calias P. 2-Hydroxypropyl-β-cyclodextrins and the Blood-Brain Barrier: Considerations for Niemann-Pick Disease Type C1 Curr Pharm Des, 2017 ; Vol. 23, pp. 6231–6238.

[40] Just N. Faber C. Probing activation-induced neurochemical changes using optogenetics combined with functional magnetic resonance spectroscopy: a feasibility study in the rat primary somatosensory cortex.J Neurochem. 2019 Aug;150(4):402–419. doi: 10.1111/jnc.14799.

